# Co-occurrence of Direct and Indirect Extracellular Electron Transfer Mechanisms during Electroactive Respiration in a Dissimilatory Sulfate Reducing Bacterium

**DOI:** 10.1101/2024.03.28.587247

**Authors:** Liyuan Hou, Rebecca Cortez, Michael Hagerman, Zhiqiang Hu, Erica L.-W. Majumder

## Abstract

Extracellular electron transfer (EET) propels microbial fuel cell (MFC) technology and contributes to the mobility of redox active minerals and microbial syntrophy in nature. Sulfate-reducing bacteria (SRB), especially the genus *Desulfovibrio* corrode metal electrodes but are of interest for sulfate-containing MFCs providing wastewater treatment. Although extensive studies on SRB-mediated metal electrode corrosion have been done, there remain knowledge gaps on SRB EET to electrodes. We aimed to determine SRB EET mechanisms towards improving SRB performance in MFC wastewater treatment. Our MFCs with *Desulfovibrio vulgaris* Hildenborough (*Dv*H), a model SRB, indicated that *Dv*H can harvest and send electrons to the carbon cloth electrode. Electricity production with a maximum power density of ∼0.074 W/m^2^ was observed when the ratio of lactate (electron and carbon donor) to sulfate (electron acceptor) was 60:20 and 0:10 in the anodic and cathodic chamber, respectively. Patterns in current production compared to variations of electron donor/acceptor ratios in the anode and cathode suggested that attachment of *Dv*H to the electrode and biofilm density were critical for effective electricity generation. Analysis of *Dv*H biofilms at different conditions (planktonic dissimilatory sulfate reduction respiration vs. electroactive respiration) by electron microscopy indicated *Dv*H utilized filaments that resemble nano-pili to attach on electrodes and facilitate EET from cell-to-cell and to the electrode. Proteomics profiling of electroactive respiration proteins indicated *Dv*H adapted to electroactive respiration by presenting more pili-, flagellar-related proteins and histidine kinases on electrodes. To investigate the role of pili and biofilm, we grew two *Dv*H mutants in MFCs under the same conditions. The mutant with a deletion of the major pilus-producing gene yielded less voltage and far less attachment to the electrode, suggesting the importance of pili in EET. The mutant with a deficiency in biofilm formation, however, did not eliminate current production indicating the existence of indirect EET. Untargeted metabolomics profiling showed flavin-based metabolites, potential electron shuttles, were dysregulated between respiration modes. This work revealed the metabolic flexibility of *Dv*H to thrive in less than ideal conditions with solid surfaces as both an electron acceptor (growth on anode) and donor (growth on cathode) by using a combination of direct and indirect EET mechanisms. Understanding *Dv*H EET mechanism could enhance the application of *Dv*H in MFCs treating wastewater.

**Importance:** We explored the application of *Desulfovibrio vulgaris* Hildenborough in microbial fuel cells (MFC) and investigated its potential extracellular electron transfer (EET) mechanism. We also conducted untargeted proteomics and metabolomics profiling, offering insights into how DvH adapts metabolically to different electron donors and acceptors. An understanding of the EET mechanism and metabolic flexibility of *Dv*H holds promise for future uses including bioremediation or enhancing efficacy in MFCs for wastewater treatment applications.

## 1. Introduction

Microbial fuel cells (MFC) have been used as a promising technology for electrical energy generation, which uses microbes to transfer the chemical energy of organic compounds into electricity (1). Novel insights have been incorporated in MFCs for energy generation as well as the microbial transformation of wastes. For instance, MFCs have been investigated for conversion of wastewater containing organic compounds and sulfate to electricity using sulfate reducing bacteria (SRB) and sulfide-oxidizing bacteria (2). These sulfate containing wastewaters are produced by many processes including mining, food processing, pulp and paper wastewater, animal husbandry, and more. (3). By culturing SRB in MFCs, previous studies have achieved the removal of sulfate and organic compounds with electricity production (4, 5), ranging from 0.013 W/m^2^ to 0.68 W/m^2^ (6–8). The overall electricity generation of an MFC mainly relies on the efficiency of extracellular electron transfer (EET) from electrogenic bacteria to the electrode (9).

Electrogenic bacteria could route their electron transport chain to the exterior of the cell through various EET mechanisms (10). Two major EET mechanisms are direct electron transfer (DET) and indirect electron transfer (IET). DET mainly relies on outer surface redox molecules and conductive nanowires. For instance, in many species, such as *Geobacter sulfurreducens*, *Shewanella oneidensis*, and *Acidithiobacillus ferrooxidans*, EET can be mediated by outer membrane *c*-type cytochromes (e.g., OmcA-MtrCAB protein complexes) (11). *G. sulfurreducens* also exchanges electrons through nanowires, which are pili formed by protein filaments (12). *S. oneidensis* MR-1 forms nanowires through extensions of the outer membrane and periplasm that include the multiheme cytochromes which are responsible for EET (13). IET involves transfer of electron through small redox active organic molecules, electron shuttles, excreted by cells or added exogenously. Different species secreted various extracellular electron carriers such as flavins and phenazine derivatives (14). Previous studies demonstrated the coexistence of DET and IET in *S. oneidensis* (15). *S. oneidensis* could simultaneously transfer electron through direct contact with the electron acceptor and also produce flavins during batch growth conditions (15). *Geobacter metallireducens* and *G. sulfurreducens* co-cultures could also secrete free redox-active flavins as the electron shuttles (16), while performing DET through cytochromes. Large numbers of studies have demonstrated that *Geobacter* and *Shewanella* species can power MFCs with high power density (11, 12, 17, 18), under conditions in proportion to their EET mechanism.

SRB can utilize organic compounds and gases (e.g., hydrogen) as electron donors (19). Recent studies also found that some SRB can use electrodes as electron donors for energy production (20). Despite this, the mechanism for SRB extracellular electron uptake is not clear due to the difficulty to distinguish the electron uptake reaction (e.g., EET mechanism) and hydrogen evolution on the electrode surface. Knowledge of EET mechanisms has major implications for being able to understand, control or intervene in several environmental problems caused by SRB, such as corrosion of steel, concrete, and electrode (21), souring of oil (22), altering mobility of toxic heavy metals (e.g., Cr and U) (23), and providing for syntrophic growth with other microorganisms (e.g., methanogens) (24). Additionally, understanding the EET mechanisms of SRB helps to improve the application and operation of SRB in MFCs.

*Desulfovibrio vulgaris* Hildenborough (*Dv*H), a model SRB strain, was reported to cause the corrosion of carbon steels due to their ability to harvest extracellular electrons from elemental iron oxidation (25). The intracellular electron transport of *Dv*H from the electron donor (i.e., lactate) to electron acceptor (i.e., sulfate) were proposed via two ways: a) hydrogen cycling pathway that uses hydrogen as an intermediate electron carrier between the periplasm and the cytoplasm, and b) a pathway that bypasses hydrogen cycling and transfers electrons directly to the membrane-bound menaquinone pool (26). However, there is little in-depth knowledge regarding the EET including DET and IET from *Dv*H to electrodes. It was found that *D. ferrophilus* IS5 was able to adopt the multi-heme cytochromes containing at least four heme-binding motifs in acquiring energy from solid electron donors (27). Kang et al. found that *D. desulfuricans* was able to conduct DET through cytochrome *c* proteins (28). However, so far, no outer membrane *c*-type cytochromes of *Dv*H have been identified (29). Deng et al. (2020) found that *Dv*H biosynthesized iron sulfide (FeS) nanoparticles on the cell membrane, which could enhance extracellular electron uptake significantly (30). Zhou et al. also demonstrated the accumulation of iron sulfide crystallite on the surface of the cell via obtaining electrons intracellularly (31). Thus, one of the DET of *Dv*H could be via iron sulfide nanoparticles. Moreover, *D. desulfuricans* utilized electrically conductive nanoscale filaments to transfer electrons to insoluble electron acceptors (i.e., iron(III) oxide) (32). However, the major characteristics of filaments of *D. desulfuricans* have not been identified. *Dv*H could also form a biofilm which is dependent on protein filaments such as flagella and pili (33). However, the role of pili, flagella and relevant biofilm in *Dv*H EET has not been investigated. Flavins, such as riboflavin and flavin adenine dinucleotide (FAD), are well-known electron shuttles (34), which carry electrons among multiple redox reactions and play an important role in IET. A recent study found riboflavin and FAD could accelerate the microbiologically influenced corrosion of 304 stainless steel by the *Dv*H biofilm (35). This indicated the potential of IET in *Dv*H. An investigation of the EET mechanism from *Dv*H to the electrode will not only fill the knowledge gap but also encourage the extensive utilizationof MFCs in the treatment of sulfate containing wastewater by SRBs.

In this study, anaerobic MFC systems adopting *Dv*H strains growing on both anode and cathode were developed. Firstly, this study aimed to determine the effects of the electron donor (i.e., lactate)/electron acceptor (i.e., sulfate) ratio in the anodic chamber and cathodic chamber, respectively, on the electricity generation. Secondly, the electricity generation of MFCs inoculated with *Dv*H JWT700, *Dv*H JW3422 (a mutant with a deletion of the gene coding for the pilin protein) and *Dv*H JWT716 (a mutant has a deficiency in biofilm formation) (36) were compared separately to unveil the role of pili and biofilm in DET of *Dv*H. Subsequently, metabolites of *Dv*H JWT700, *Dv*H JW3422, and *Dv*H JWT716 under electroactive respiration were analyzed to screen potential electron shuttles. At last, the conductivity of the surface structure of *Dv*H and their role in electricity generation were evaluated.

## 2. Materials and methods

### 2.1 Bacteria strains and culture cultivation

All *Desulfovibrio vulgaris* Hildenborough strains were isogenic of wild-type strain JWT700. A deletion mutant *Dv*H JWT716 was used in this study, which lacks the type 1 secretion system’s ABC transporter protein resulting in a deficiency in biofilm formation (36). The deletion mutant *Dv*H JW3422 strain, which is a deletion of the gene coding for the pilin protein (DVU2116), was constructed using the same marker replacement plasmid and method as strain JW9003 from a previous study (37). The plasmid for deleting DVU2116 and replacing with a kanamycin resistance gene was transformed into *Dv*H JWT700 and deletion was confirmed by Southern Blot.

*Dv*H strains were grown anaerobically in routine for 15 h at 34 °C on lactate (60 mM), sulfate (30 mM), and a nutrient medium, named as MOYLS4 (MgCl_2_, 8 mM; NH_4_Cl, 20 mM; CaCl_2_, 0.6 mM; NaH_2_PO_4_·H_2_O, 2 mM; FeCl_2_, 0.06 mM; EDTA, 0.12 mM; Thioglycolate, 1.2 mM; 6 ml of trace elements solution per liter; and 1 ml of Thauer’s vitamin solution per liter) with the pH adjusted to 7.2. The trace elements solution contained 0.5 g/L MnCl_2_, 0.3 g/L CoCl_2_, 0.2 g/L ZnCl_2_, 0.05 g/L Na_2_MoO_4_, 0.02 g/L H_3_BO_3_, 0.1 g/L NiSO_4_, 0.002 g/L CuCl_2_, 0.006 g/L Na_2_SeO_3_, and 0.008 g/L Na_2_WO_4_. Thauer’s vitamin solution contained 2 mg of biotin, 2 mg of folic acid, 10 mg of pyridoxine HCl, 5 mg of thiamine HCl, 5 mg of riboflavin, 5 mg of nicotinic acid, 5 mg of D calcium pantothenate, 0.1 mg of vitamin B12, 5 mg of *p*-amionobezoic acid and 5 mg of lipoic acid, in 1 L of deionized water (38). These media were flushed with nitrogen gas for 15 min in Balch anaerobic tubes sealed with a butyl rubber stopper before use. In this medium, the metabolism of *Dv*H results in the production of H_2_S, which reacts with Fe and generates FeS (black color). The anaerobic tubes were inoculated with a log phase culture of *Dv*H to an optical density of 0.8 at 600 nm (OD_600_). The bacterial absorbance was measured at 600 nm using a spectrophotometer (GENESYS 10S UV-VIS, Thermo Scientific, USA).

### 2.2 MFC construction and operation

Double-chambered MFC were fabricated with a working volume of approximately 90 mL for each compartment and were used throughout the study (Fig. 1). Both anode and cathode were made of carbon cloths (60 wt % Vulcan XC-72 and loaded with Pt 0.5 mg/cm^2^, Fuel Cell Store, USA). Nafion N117 (Fuel Cell Store, USA) was used as a proton exchange membrane (PEM) in the system and pretreated as it was described in the literature (39). The electrodes were connected using platinum wire. The distance between the anode and the cathode was approximately 10 cm. The MFC was operated under a constant external resistance of 1 kΩ using a pure culture of *Dv*H cells growing in both the anodic and cathodic chamber. The bottles and accessories were autoclaved or sterilized before they were assembled.

**Figure 1.**
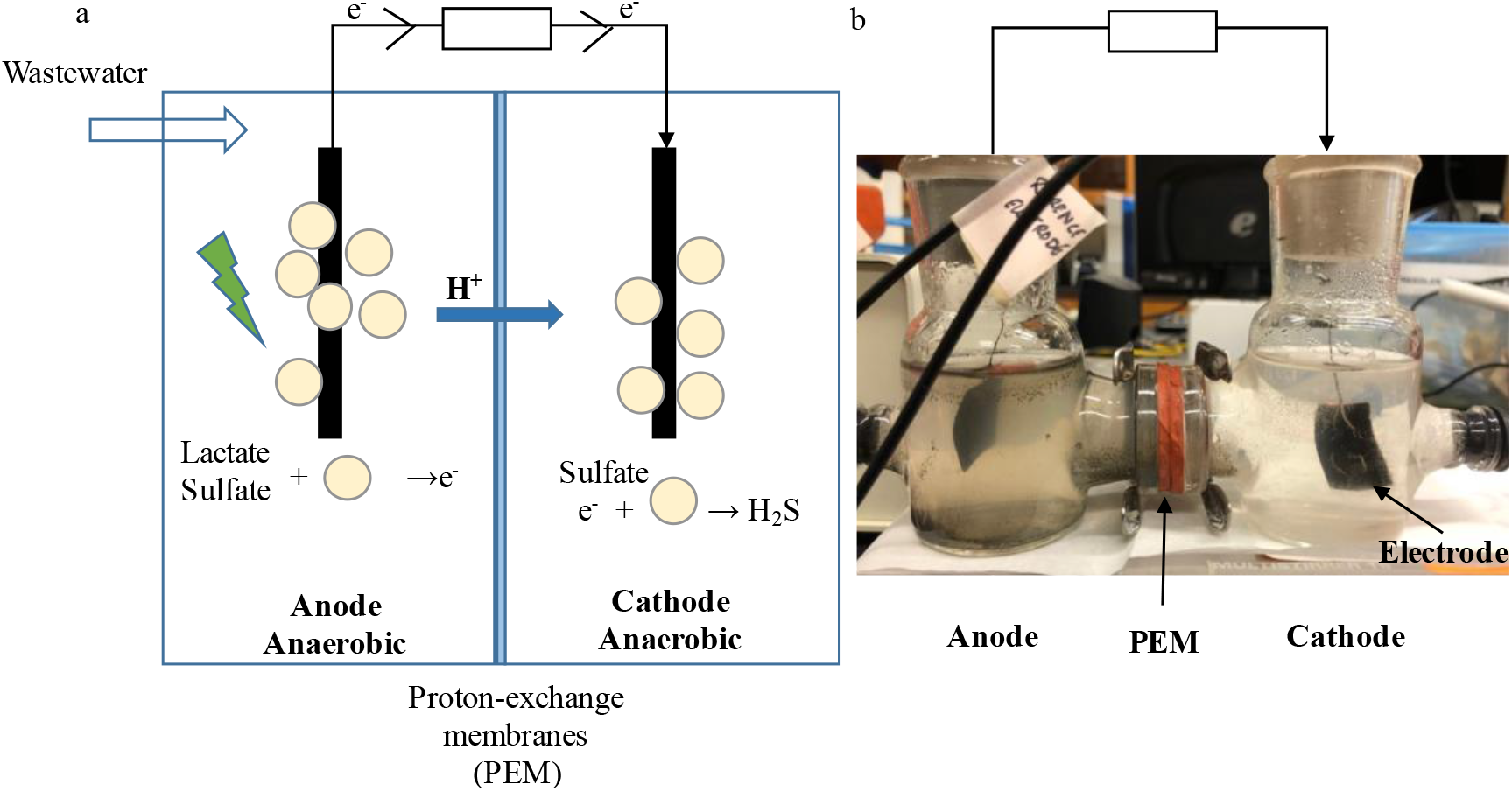
Schematic (a) and laboratory-scale prototype (b) of the MFC.

The anodic chamber and cathodic chamber of the MFC were inoculated with modified MOYLS4 media that contained different concentrations of lactate and sulfate as was shown in Table 1. The anodic chamber was fed with 60 mM lactate while 0 mM lactate was provided in the cathodic chamber. Both anodic and cathodic chamber was operated an initial pH of 7.2 in all experimental runs. All modified MOYLS4 media for each run were autoclaved at 121 °C for 20 min. The anode and cathode were kept anaerobic by sparing filter-sterilized nitrogen gas for 15 min at the beginning of each run. In each run, *Dv*H seed culture with an optical density (OD_600_) of 0.8 was used to inoculate each anodic and cathodic chamber to achieve an initial cell concentration of approximately 10^7^ cells/mL right after inoculation. Agitation was maintained at 50 rpm to minimize mechanical shear forces.

**Table 1.** MFC experimental conditions.

| Experiment<br>al run | Strain | Size of<br>anode<br>(cm × cm) | Lactate in<br>anode<br>chamber<br>(mM) | Sulfate in<br>anode<br>chamber<br>(mM) | Lactate in<br>cathode<br>chamber<br>(mM) | Sulfate in<br>cathode<br>chamber<br>(mM) |
| --- | --- | --- | --- | --- | --- | --- |
| Control | *DvH*<br>JWT700 | 2 × 2 | 0 | 0 | 0 | 30 |
| Run 1 |  | 2 × 2 | 60 | 0 | 0 | 10 |
| Run 2 |  | 2 × 2 |  | 10 |  | 10 |
| Run 3 |  | 2 × 2 |  | 20 |  | 10 |
| Run 4 |  | 2 × 2 |  | 30 |  | 10 |
| Run 5 |  | 2 × 2 |  | 10 |  | 0 |
| Run 6 |  | 2 × 2 |  | 10 |  | 20 |
| Run 7 |  | 2 × 2 |  | 10 |  | 30 |
| Run 8 |  | 2 × 2 |  | 20 |  | 30 |
| Run 9 |  | 3 × 3 |  | 20 |  | 30 |
| Run 10 | *DvH*<br>JW3422 | 2 × 2 |  | 20 |  | 30 |
| Run 11 | *DvH*<br>JWT716 | 2 × 2 |  | 20 |  | 30 |

Run 1 to run 7 was designed to investigate voltage generation at different sulfate concentrations in the anodic chamber and cathodic chamber, respectively (Table 1). For run 1 to run 7, *Dv*H JWT700 was applied, and the electrode size was 2 cm × 2 cm. A control experiment (run 0) with 30 mM Na_2_S adding in the anodic chamber to investigate the role of S^2-^ in the electron transfer with *Dv*H was conducted. Optimal condition (the lactate and sulfate concentration in each chamber) for potential maximum voltage production based on the results of run 1 to run 7 was proposed, where the ratio of lactate to sulfate is 60:20 in the anodic chamber and 0:30 in the cathodic chamber. Run 8, 9, 10, and 11 were operated under this condition. To investigate the effect of anode size on voltage generation, run 9 adopted a larger anode size (3 cm × 3 cm) compared to run 8 (2 cm × 2 cm). *Dv*H JW3422 and JWT716 strain were applied with an anode and cathode size of 2 cm × 2 cm to test the voltage production without pili and biofilms in run 10 and run 11, respectively. All experiment runs were incubated at room temperature (25 ± 1 °C) throughout the operation. Duplicate experiments were conducted for each run. Voltages across the electrodes were measured over time during MFC operation until the voltages decreased to zero. Samples were obtained at the end of each operation to measure pH, acetate, sulfate concentration after filtration (0.22 μm nylon syringe filter).

### 2.3 Chemical analyses and electricity generation calculations

Acetate concentrations were quantified by HPLC (Shimadzu LC-20A) equipped with a Supelcogel C610H (30 cm × 7.8 mm) column (Supelcogel, PA) (40). The mobile phase was 0.1% H_3_PO_4_ and HPLC samples were measured at a wavelength of 210 nm. Sulfate concentrations were determined spectrophotometrically (41). Voltage was carried out in the open circuit potential time mode using a CHI600 (Austin, TX, USA) electrochemical workstation. The current was calculated using Ohm’s law, *I* = *V*/*R*, where *I* = current (mA), *V* = voltage (mV), *R* = resistance (Ω). The power density (W/m^2^) was normalized by the surface area of the anodic chamber. The coulombic efficiency (C_E_) was calculated using the following equation: *C_E_* = *C*_P_/*C*_T_ × 100%, where *C*_P_ is the harvested coulombs calculated by integrating the current over operation time, and *C*_T_ is the theoretical number of coulombs that can be produced from the substrate used. *C_T_* was calculated using the formula: *C_T_*= *F*nSv, where *F* is Faraday’s constant (96,487 C mol^−1^ electron), n is the mol number of electrons produced per mol of substrate oxidation (n = 4), S is the substrate (lactate) concentration (mol), v is the effective volume of the MFC (L). The energy efficiency was calculated as *η*_E_= *E_p_*/*E_T_*×100 %, where *E_p_* is the harvested energy (in joules) calculated by integrating the power (P = I × V) over operational time and *E_T_* is the theoretical value of available energy, obtained from the change in Gibbs free energy, Δ*G*, of - 160.1 kJ mol^−1^ lactate oxidation with sulfate as the electron acceptor (42).

### 2.4 Electrochemical analysis

Cyclic voltammetry (CV) was employed to study the electrochemical performance of electrodes and media with various *Dv*H mutants. Anodes and cathodes from stable running MFCs (run 8, run 10, and run 11) were thoroughly rinsed with phosphate buffered saline (pH 7.4, 137 mM NaCl, 2.7 mM KCl, 8 mM Na_2_HPO_4_, and 2 mM KH_2_PO_4_) and assembled as working electrodes in a three-electrode cell with Pt wire as counter electrode, Ag/AgCl (sat. KCl, 222 mV *vs.* SHE) electrode as the reference electrode, and phosphate buffered saline as supporting electrolyte. The reference electrode was disinfected with 75% ethanol and was then placed in the vicinity of the working electrode surface during CV measurements. The scan rate was 10 mV/s over a range from −0.8 to +0.6 V vs. Ag/AgCl. Electrochemical impedance spectroscopy (EIS) was performed to elucidate the resistance characteristics for various anodes and cathodes with AC signal amplitude of 10 mV. Charge transfer resistances were obtained by external equivalent circuit fitting analysis of the EIS data, using the Randle’s circuit. For the culture media in anodic and cathodic chamber, 50 mL suspension from each chamber in run 8, 10, and 11 were collected to be tested by CVs with glassy carbon as the working electrode, Ag/AgCl (sat. KCl, 222 mV *vs.* SHE) electrode as the reference electrode and Pt wire as the counter electrode. The CV scan rate was 10 mV/s ranging from −0.6 V to +0.7 V vs. Ag/AgCl.

### 2.5 Scanning electron microscopy (SEM)

The electrodes from both the anodic and cathodic chamber were obtained from run 8, run 10, and run 11, respectively. A 5 mm × 5 mm section of carbon cloth was cut off from each electrode for SEM analysis. The fixation procedure was done as described elsewhere (32). Briefly, electrode samples were placed in a fixative that contained 2.5% glutaraldehyde (w/v), 2.0% paraformaldehyde (w/v) and 0.05 mM sodium cacodylate buffer (pH 7.0). The biofilms were fixed overnight and were then washed four times with double-distilled water. Electrode samples were dehydrated by incubation in increasing concentrations of ethanol and then dried. A FEI Quanta 600 FEG SEM was used to examine the surfaces of the biofilm formation on the electrode surfaces. Along with the SEM, X-ray energy dispersive spectroscopy (EDS) mapping was employed to detect C, O, N, S, F, and Fe elements on the surface of Pt treated conversion coating.

### 2.6 Conductivity measurements of the filament

A size of 10 mm × 10 mm carbon cloth obtaining from run 8 was cut off from the anode and cathode, respectively. Electrode samples were pretreated using the same fixation procedure described above. Topography images were measured in ambient conditions using tapping mode atomic force microscopy. The local conductivity was completed using the contact mode of a Veeco Dimension V scanning probe microscope. Morphology characterization was completed in tapping mode using Veeco RTESP or OTESPA probes. For stable current-voltage characterization (CVC) responses, applied voltage bias was 2 V and 4 V, respectively, for each electrode sample. The maximum applied bias of the module was 10 V. The CVC response of the plain carbon cloth was measured as well as the control.

### 2.7 Protein concentration on electrodes and in suspensions

MFC setups that cultivated with *Dv*H JWT700, *Dv*H JW3422, and *Dv*H JWT716 (i.e., run 8, run 10, and run 11), respectively, were repeated and operated for five days. After five days, 10 mL solution samples were obtained from each chamber as the suspension sample for protein concentration measurement and peptide sequence analysis. To measure the total protein concentration on the electrodes, the anode and cathode were immersed in 10 mL lysis buffer, which contains 50 mM Tris-HCl and 200 mM NaCl. Then, all these samples were sonicated on ice for 5 min (with 30-second intervals to prevent overheating between each 1-minute cycle) and centrifuged at 8,000 × g for 20 min to obtain the supernatants, which were used to measure the protein concentrations. A portion of each of the samples from run 8, 10 and 11 was stored in ‒ 80°C for peptide sequence analysis later. The total protein concentration was measured using the Bradford protein assay and the absorbance of samples was measured at 595 nm. Bovine serum albumin (BSA) was used as standard and measured in a concentration range of 5∼100 µg/mL along with the analysis of each unknown sample.

### 2.8 Proteomics

#### 2.8.1 Enzymatic “In Liquid” Digestion, TMT labelling and high pH separation

*Desulfovibrio vulgaris* Hildenborough cell suspensions (1 mL) were transferred to 2ml microcentrifuge tube, 125 µL of protease inhibitors were added immediately (10x cOmplete Mini from Roche) along with 125ul of 10% SDS. Cell lysis was conducted by probe sonication with 2×20sec intervals at 3W setting with cooling on ice in between. Samples were subsequently spun for 4 minutes at max speed (room temperature) to pellet cellular debris and supernatant (1,250 µL) was transferred to new 2ml tube. Protein precipitation was initiated next by 200 µL addition of 100% trichloroacetic acid and 550ul of cold acetone, samples were incubated on ice for 1 hour the spun for 10 minutes at max speed (room temperature). Generated protein pellets were washed twice with cold acetone, once with cold 80% methanol then finally with 100% cold methanol and air dried. Pellets were solubilized in 8 M Urea in 25 mM NH_4_HCO_3_ (pH 8.5), 5 µL aliquot was taken for BCA protein measurement and the rest was used for tryptic/LysC digestion where the samples were first reduced with 1mM DTT for 15 minutes at 56°C. After cooling on ice to room temperature 2.6 mM CAA (chloroacetamide) was used for alkylation where samples were incubated in darkness at room temperature for 15 minutes. This reaction was quenched with 3.7mM DTT. Subsequently Trypsin/LysC solution [100 ng/μL 1:1 Trypsin (Promega):LysC (FujiFilm) mix in 25mM NH_4_HCO_3_] was added for ∼1:40 enzyme:substrate ratio and 25 mM NH_4_HCO_3_ (pH 8.5) was added to the samples for a final 100 µL volume. Digests were carried out overnight at 37°C then subsequently terminated by acidification with 2.5% TFA [Trifluoroacetic Acid] to 0.3% final. 10% HFBA [Heptafluorobutyric acid] was added to 0.2% final and each individual sample was cleaned up using 100 µL Bond Elut OMIX C18 pipette tips (Agilent) according to manufacturer protocol. Eluates in 70%:30%:0.1% acetonitrile:water:TFA acid (vol:vol) were dried to completion in the speed-vac and reconstituted in 25 µL (Low-abundance), 50ul (Medium-abundance) or 100 µL (High-abundance) of 100 mM TEAB [Triethylammonium Bicarbonate] for TMTpro labelling with 5 µl for Low-abundance, 10 µL for Medium-abundance and 20 µl for High-abundance of TMTpro reagent [TMTpro 16plex™ Labeling Reagent Set Lot#XE350091 at 12.5 µg/µL from Thermo Scientific] done at room temperature for 1 hr with intermittent gentle vortexing. Digested Bovine Serum Albumin internal protein standard was added during labeling to each sample for downstream normalization (2 µg for High, 1 µg for Medium and 500 ng for Low set). Additionally, unused TMT channels per each set were used to label a pooled control sample (even aliquot of each individual sample) to be used as a possible normalization standard between independent sets. The labeling reaction was terminated with addition of 5% hydroxylamine (0.2% final) and 15-minute incubation at room temperature. Master-pool samples were generated by combining all of the individual labeled reactions of each set, freezing at −80⁰C and drying to completion using speed-vac. Subsequently re-solubilized with 0.3% TFA / 0.2% HFBA and solid-phase extracted with Pierce C18 spin tips (Low-set in 200ul containing ∼68 µg total digested protein) or Phenomenex Strata-X 33um polymeric column [10mg/1ml size for Medium-set in 400 µL containing ∼380 µg total digested protein or 30 mg/3 mL size for High-set in 800 µL containing ∼860 µg total digested protein] according to manufacturer protocol. Eluates were dried and finally reconstituted in 0.1% Formic acid / 3% Acetonitrile to 1.4 µg/µL concentration for each set.

#### 2.8.2 NanoLC-MS/MS

Peptides were analyzed by Orbitrap Fusion™ Lumos™ Tribrid™ platform, where 2 µL was injected using Dionex UltiMate™3000 RSLCnano delivery system (ThermoFisher Scientific) equipped with an EASY-Spray™ electrospray source (held at constant 50°C). Chromatography of peptides prior to mass spectral analysis was accomplished using capillary emitter column (PepMap® C18, 2µM, 100Å, 500 x 0.075mm, Thermo Fisher Scientific). NanoHPLC system delivered solvents A: 0.1% (v/v) formic acid, and B: 80% (v/v) acetonitrile, 0.1% (v/v) formic acid at 0.30 µL/min to load the peptides at 2% (v/v) B, followed by quick 2 minute gradient to 5% (v/v) B and gradual analytical gradient from 5% (v/v) B to 62.5% (v/v) B over 203 minutes when it concluded with rapid 10 minute ramp to 95% (v/v) B for a 9 minute flash-out. As peptides eluted from the HPLC-column/electrospray source survey MS scans were acquired in the Orbitrap with a resolution of 60,000 followed by HCD-type MS2 fragmentation into Orbitrap (36% collision energy and 30,000 resolution) with 0.7 m/z isolation window in the quadrupole under ddMSnScan 1 second cycle time mode with peptides detected in the MS1 scan from 400 to 1400 m/z; redundancy was limited by dynamic exclusion and MIPS filter mode ON.

#### 2.8.3 *Proteomics Data* analysis

Raw data was directly imported into Proteome Discoverer 2.5.0.400 where protein identifications and quantitative reporting was generated. Seaquest HT search engine platform was used to interrogate Uniprot *Desulfoovibrio vulgaris* reference proteome database (UP000002194, 09/28/2022 download, 3,519 total entries) along with a cRAP common lab contaminant database (116 total entries). Cysteine carbamidomethylation and TMTpro specific labeling was selected as static modifications whereas methionine oxidation and asparagine/glutamine deamidation were selected as dynamic modifications. Peptide mass tolerances were set at 10ppm for MS1 and 0.02Da for MS2. Peptide and protein identifications were accepted under strict 1% FDR cut offs with high confidence XCorr thresholds of 1.9 for z=2 and 2.3 for z=3. For the total protein quantification processing Reporter Ion Quantifier settings were used on unique and razor peptides, protein grouping was considered for uniqueness. Reporter abundance was based on BSA-normalized peptide amount intensity values, scaled on all average and with co-isolation threshold filter set at ≤40. ANOVA (individual proteins) hypothesis was executed without imputation mode being executed.

### 2.9 Untargeted metabolomics using high-resolution LC-MS

A total of 20 mL suspension samples in both chambers was obtained separately from run 8, run 10, and run 11, respectively. Samples were centrifuged at 4000 rpm for 10 min and the supernatant (∼15 mL) was transferred to a new tube for metabolites extraction. Metabolites from 5 mL samples were extracted by three freeze-thaw cycles with liquid nitrogen following by sonication in the same volume of a 2:2:1 acetonitrile-methanol -water mixture. Samples were then placed in −20°C overnight to allow proteins and cell debris to precipitate. Then, samples were centrifuged for 15 min at 13,000 rpm and 4°C. Supernatants were transferred to high recovery glass vials and dried in the speed vac at 10 °C. Dried metabolite extracts were reconstituted in volumes of acetonitrile: water (1:1, v/v) normalized to protein content in the sample as determined using NanoDrop Protein A280 measurement mode. The resulting metabolite suspension was dried down in a speed-vac overnight. Metabolites were resuspended in 1:1 acetonitrile: water.

Mass spectrometry data were acquired by running a Thermo Scientific Q Exactive HF Orbitrap LC-MS/MS system in both positive and negative ion modes. Accucore™ Vanquish™ C18 column (2.1 × 100 mm, 1.5 μm, Thermo Scientific) and Accucore™ 150-Amide-HILIC (4.6 × 100 mm, 2.6 μm, Thermo Scientific) were used in the separation of metabolites in positive and negative modes, respectively, with a 5 μL injection volume. Full MS-ddMS^2^ detection mode was applied. For LC, the mobile phases comprised water containing 0.1% formic acid (A) and acetonitrile containing 0.1% formic acid (B). For the reverse phase analysis, metabolites were separated by gradient elution at a flow rate of 0.25 mL/min starting at 5% (v/v) B, held for 5 min, increased to 99.5% B within 20 min, and reverted to 5% B at the 30th min, held for 2 min, with a total run time of 32 min. For the HILIC analysis, metabolites were separated by gradient elution at a flow rate of 0.5 mL/min starting at 1% (v/v) A, increased to 35% A within 15 min, then to 60% A at the 18th min, and reverted to 1% A at the 20th min, with a total run time of 20 min.

The mass spectrometer was operated as follows: spray voltage 3.5 kV, and capillary temperature 350°C. The flow rates of sheath gas, aux gas and sweep gas were set to 50, 2, and 0, respectively. Full MS resolution was set to 120,000, full MS AGC target was 3×10^-6^ with a maximum IT of 250 ms. The scan range was set to 100∼1000 m/z. For MS2 spectra, the AGC target value was set to 2×10^-5^, isolation width was set to 1 m/z. The resolution was set to 15,000 and normalized collisional energy was 40. The dynamic exclusion duration was set to 1.5 s.

Raw data files were converted and then processed using XCMS Online (45, 46) as a multigroup job comparing under different electroactive respiration conditions. For each sample, the filtered features data-table was annotated via an accurate mass search against METLIN (47).

### 2.10 Data Availability Statement

All data are provided within this manuscript. Raw proteomics and metabolomics files as well as strains will be made available upon request.

## 3 Results

### 3.1 Electricity generation in the MFC under different sulfate concentrations

To examine the feasibility and optimal conditions for electricity generation by using *Dv*H JWT700 in the MFC, different sulfate (electron acceptor) concentrations were tested in two chambers, where a fixed concentration of lactate (electron donor; 60 mM) was applied to the anodic chamber while no lactate was provided in the cathodic chamber. The half reaction on the anode is listed as follows: C_3_H_5_O_3_^-^ + H_2_O → C_2_H_3_O_2_^-^+4e^-^+4H^+^+CO_2_. On the cathode, the reduction of sulfate occurs: SO_4_^2-^+ 10H^+^+ 8e^-^→ H_2_S+ 4H_2_O. According to the stoichiometry coefficient, run 1, run 2, run 3, run 4, run 5 and run 6 can theoretically utilize 33.3%, 66.7%, 100%, 100%, 33.3%, 100%, and 100% of the lactate. Among different treatments, electricity generations were observed for run 1, run 2 and run 3, where the maximum electricity generation increased with the increase of the sulfate concentration (Fig. 2a). However, as for run 4, no significant electricity generation was observed. Moreover, when the concentration of sulfate in the cathodic chamber increased and the condition in the anodic chamber stayed the same, a higher maximum electricity generation occurred, and the duration of electricity generation extended (Fig. 2b). A longer lag period was observed for the treatments with a higher maximum electricity generation (Fig. 2a and b).

**Figure 2.**
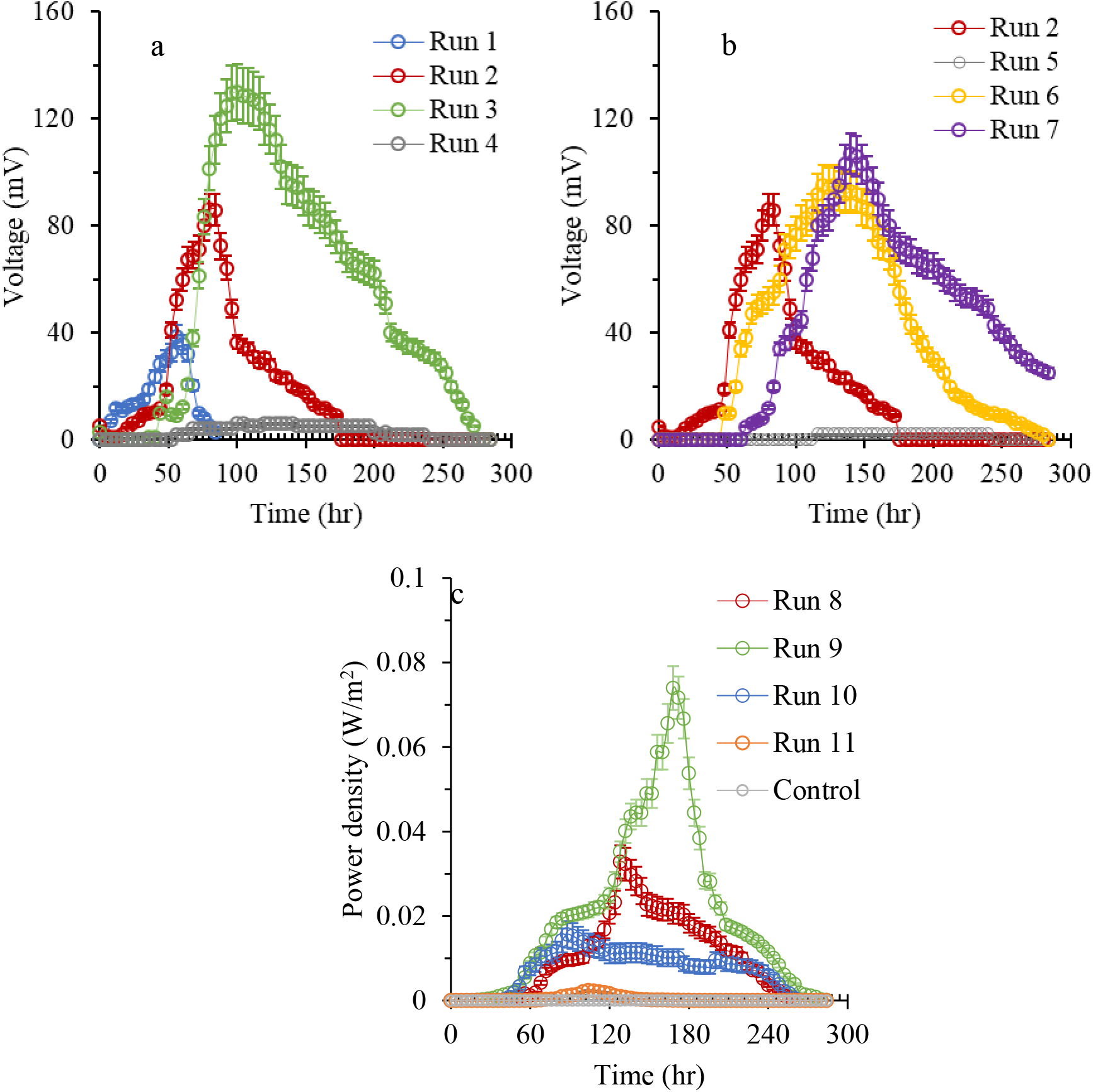
Effects of lactate/sulfate ratio to the anode chamber (a) and cathode chamber (b) on electricity generation. (c) Effect of electrode size and different *D. vulgaris* Hildenborough stains in the anode chamber on powder density.

Among these treatments, a maximum voltage of 131 mV was achieved in run 3 (Table 2). The coulombic efficiencies ranged from 0.79% to 3.13%. The corresponding energy efficiency varied from 0.01% to 0.69%. A slight increase in pH was observed at the end of each test in the cathodic chamber. Thus, the condition with a sulfate concentration of 20 mM, 30 mM in the anodic and cathodic chamber, respectively, was proposed for a higher electricity generation (Table 1).

**Table 2.**
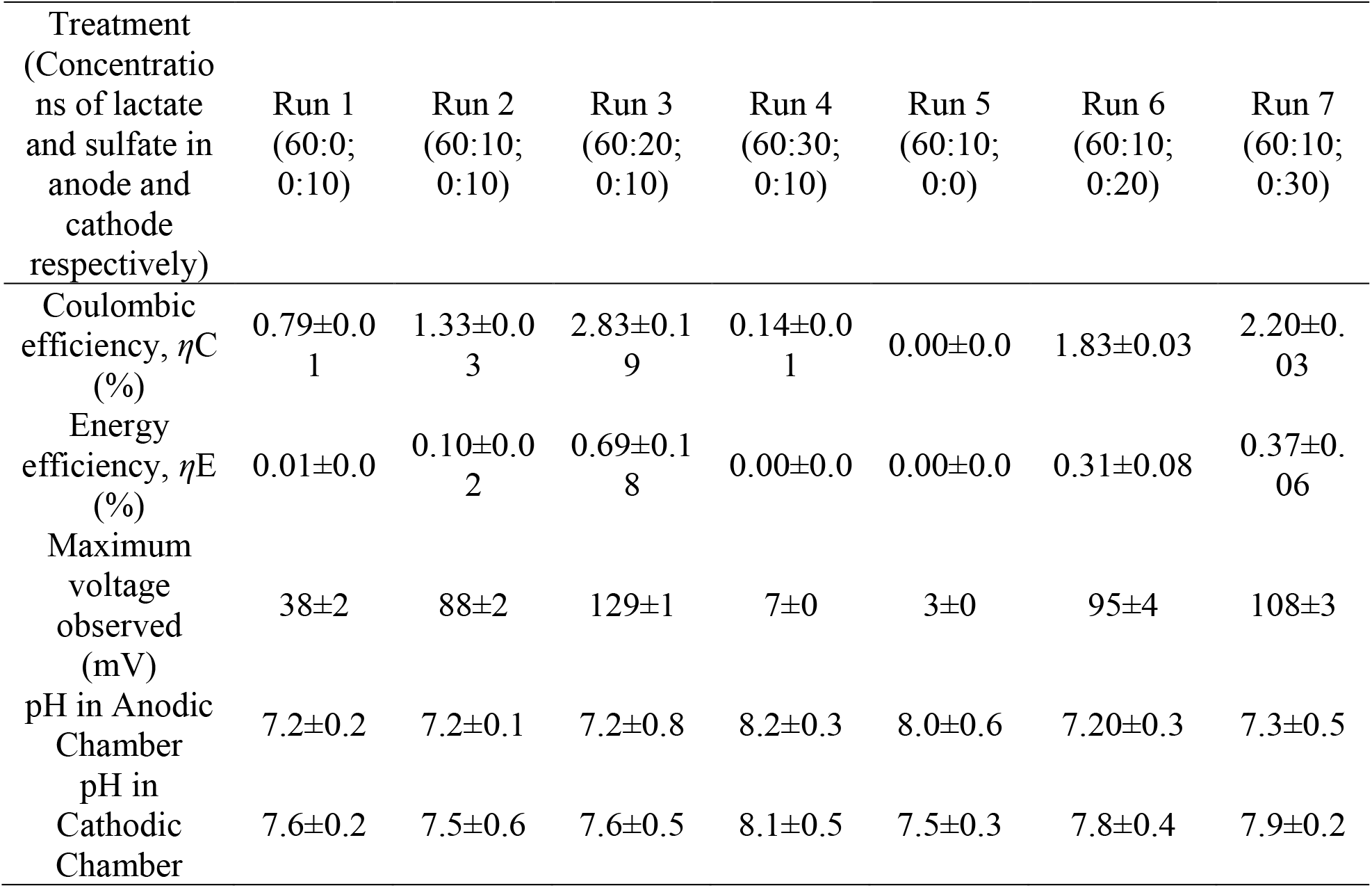
Comparison of electricity generation, coulombic efficiencies, energy efficiency and pH in MFCs using *Dv*H JWT700 under different treatments.

In the anode chamber, *Dv*H JWT700 converted 1 mol of lactate to 1 mol of acetate, producing four electrons. The theoretical electron generation in the anode chamber of run 1, run 2, run 3, and run 4 is 0, 80, 160 and 240 e^-^ eq/L, respectively, according to the initial sulfate concentration in the anodic chamber. However, a higher electron generation was observed for these operations except run 4 based on the acetate production, which are 14.2 ± 2.1, 107.9 ± 5.4, 192 ± 6.9, 242 ± 8.8 e^-^ eq/L (Fig. S1a). As the lactate/sulfate ratio increased, additional acetate was produced except the ratio value of 2. In the cathodic chamber, 1 mol of sulfate needs eight electrons to be reduced into sulfide. The electron required in the cathode chamber of run 1, run 2, run 3, and run 4 is 8 ± 1.7, 25.2 ± 4.2, 31.8 ± 5.6 and 1.1 ± 0.7 e^-^ eq/L, respectively, according to the residual sulfate concentration in cathodic chamber (Fig. S1b). This could be due to the electron transfer from anode to cathode.

Similarly, the theoretical electron generation in the anode chamber of run 2, run 5, run 6, and run 7 is the same, which is 80 e^-^ eq/L based on the initial sulfate concentration in the anodic chamber. However, the actual electron generations were 107.6 ± 6.8, 81.3 ± 2.1, 123.2 ± 3.4, 147.6 ± 5.6 e^-^ eq/L for run 2, 5, 6 and 7, respectively (Fig. S1c). As the sulfate ratio increased in the cathodic chamber, more extra acetate was generated. Correspondingly, 25.2 ± 3.6, 0.0 ± 0.0, 34.6 ± 2.8 and 66.3 ± 4.3 e^-^ eq/L were required in the cathode chamber for run 2, 5, 6 and 7, respectively (Fig. S1d). This indicated that most of the electrons generated from the anode were transferred to the cathode for sulfate reduction. In addition, Run 5, which exhibits no electron transfer, is attributed to the sulfate in the anodic chamber capturing all electrons generated by dissimilatory sulfate reduction in *Dv*H with lactate as the electron donor.

### 3.2 Effects of different electrode sizes and *Dv*H mutants on electricity generation

Power densities of MFCs using different anode sizes and *Dv*H strains in MFCs were shown in Fig. 2c. All three MFC set-ups were run for around 12 days until the electricity generation approached zero. It was observed that after a lag phase of 28 h, the electricity started to be produced and gradually declined after 130 h. Increasing the electrode size, indicating a larger surface area for microbes to transfer electrons, may result in the increase of the electricity generation for MFCs. As expected, the anode with a larger surface area (run 9; 3 cm × 3 cm) produced a higher electric potential compared to the anode with a surface area of 2 cm × 2 cm (run 8) (Table 3). Run 9 exhibited a maximum power density of ∼0.074 W/m^2^, indicating that the larger electrode size may not only offer more significant surface areas for *Dv*H but also enhance its attachment. Run 10 applying the *Dv*H JW3422, which could not form pili, had a lower electrical output of ∼0.015 W/m^2^ compared to run 8 (∼0.040 W/m^2^) (Fig. 2c). This suggested that the pili formed by *Dv*H JWT700 may contribute to electricity generation. In addition, there is no significant electricity generation output for *Dv*H JWT716, which has a deficiency in biofilm formation in run 11 (Fig. 2c).

**Table 3.** Comparison of power generation, coulombic efficiencies, and pH in MFCs with different electrode sizes and different *Dv*H stains in anode chamber.

| Treatment | Run 8<br>(*DvH*<br>JWT700; 2 ×<br>2 cm<sup>2</sup>) | Run 9<br>(*DvH*<br>JWT700; 3 ×<br>3 cm<sup>2</sup>) | Run 10<br>(*DvH*<br>JW3422; 2 ×<br>2 cm<sup>2</sup>) | Run 11<br>(*DvH*<br>JWT716; 2<br>× 2 cm<sup>2</sup>) |
| --- | --- | --- | --- | --- |
| Coulombic efficiency, $\eta_C$ (%) | 2.65±0.01 | 5.65±0.05 | 2.52±0.06 | 0.36±0.02 |
| Energy efficiency, $\eta_E$ (%) | 0.56±0.01 | 2.12±0.06 | 0.42±0.02 | 0.02±0.01 |
| Maximum voltage observed (mV) | 125±1 | 248±2 | 87±3 | 42±4 |
| pH in Anode Chamber | 7.2±0.2 | 7.3±0.1 | 7.5±0.2 | 7.2±0.2 |
| pH in Cathode Chamber | 7.6±0.3 | 7.7±0.2 | 7.7±0.3 | 7.4±0.3 |

**Table 4.** Protein concentration on electrodes and solutions for MFC cultivating different DvH strains.

|  | Anodic chamber |  | Cathodic chamber |  |
| --- | --- | --- | --- | --- |
|  | Anode<br>Total protein<br>(µg) | Solution<br>Protein concentration<br>(µg/ml) | Cathode<br>Total protein<br>(µg) | Solution<br>Protein concentration<br>(µg/ml) |
| Initial setup | 0 | 3.9 ± 2.3 | 0 | 3.9 ± 2.3 |
| DvH JWT700<br>(Run 8) | 262.2 ± 6.2 | 54.9 ± 0.7 | 8.5 ± 3.1 | 12.7 ± 0.9 |
| DvH JW3422<br>(Run 10) | 186.3 ± 19.6 | 56.7 ± 1.2 | 1.1 ± 0.5 | 10.7 ± 0.5 |
| DvH JWT716<br>(Run 11) | 27.7 ± 3.1 | 42.0 ± 3.5 | 0.1 ± 0.0 | 5.9 ± 0.6 |

According to the initial sulfate concentration in the anodic chamber, the theoretical electron generation in the anode chamber of run 8, 9, 10, and 11 are the same, which is 160 e^-^ eq/L, However, run 9 with a larger electrode size had a significantly higher electron generation (238.2 ± 6.3 e^-^ eq/L) than run 8 (224.0 ± 3.1 e^-^ eq/L) (Fig. S2a). As such, more sulfate was reduced in run 9 than that in run 8 in the cathodic chamber (Fig. S2b). Additionally, run 10 with non-pili forming strain *Dv*H JW3422 had an electron generation of 189.6 ± 5.4 e^-^ eq/L in anode chamber while run 10 with non-biofilm forming strain *Dv*H JWT716 produced 161.0 ± 2.3 e^-^ eq/L. Correspondingly, 2.83 ± 0.08 mM and 0.24 ± 0.01 mM sulfate were reduced by using 22.7 ± 0.7 and 1.9 ± 0.08 e^-^ eq/L sending from anode to cathode, for run 10 and 11, respectively. Table 3 also showed that pH in the cathodic chambers significantly increased after different treatments (*p* < 0.01). By conducting a control experiment using Na_2_S to mimic the S^2-^ produced in the anode chamber, no electricity signal was observed. This indicated that S^2-^ was not the main factor for electricity generation.

The initial protein concentration after seeding in the solution in both chambers of MFCs was 3.9 ± 2.3 µg/mL. After ∼5 days of operation, the protein concentrations in the anodic chamber increased due to the growth of cells. It was observed that MFCs with *Dv*H JWT700 and *Dv*H JW3422 had higher protein concentrations than the MFC with *Dv*H JWT716 in the anodic chamber. Additionally, the protein concentrations in the cathodic chamber for MFCs with *Dv*H JWT700 and *Dv*H JW3422 increased from 3.9 ± 2.3 µg/mL to 12.7 ± 0.9 µg/mL and 10.7 ± 0.5 µg/mL, respectively. The anode with *Dv*H JWT700 had the highest amount of the total protein (262.2 ± 6.2 µg), followed by the anode with *Dv*H JW3422 (186.3 ± 19.6 µg) and then the anode with *Dv*H JWT716 (27.7 ± 3.1 µg). A similar trend was found for the protein amount on the cathodes with various *Dv*H strains: *Dv*H JWT700 (8.5 ± 3.1 µg) > *Dv*H JW3422 (1.1 ± 0.5 µg)> *Dv*H JWT716 (0.1 ± 0.0 µg). However, the amounts of protein on the cathode were much lower than the ones on the anode.

### 3.3 Electrochemical analysis of electrodes and electrolytes with different *Dv*H mutants

CVs were collected using the anodes and cathodes with biofilms formed by *Dv*H JWT700, *Dv*H JW3422, and *Dv*H JWT716, respectively, as working electrodes to determine the effects of *Dv*H mutants on electrodes’ performance (Fig. 3). Compared to electrodes with *Dv*H mutants, the plain electrode (*i.e.,* carbon cloths) displayed greater current densities and larger CV curve areas without obvious distortions, demonstrating its higher capacitive behaviors. Electrodes with *Dv*H mutants presented a well-defined symmetric shape and similar enclosed area, suggesting a comparable electric double layer (EDL) capacitive performance. In particular, anodes with *Dv*H JW3422 had a remarkable oxidation peak at about −0.05 V vs. Ag/AgCl, while the redox reactions presented the irreversibility without the presence of mirrored reduction peak. A similar trend was not observed for the cathode with *Dv*H JW3422.

**Figure 3.**
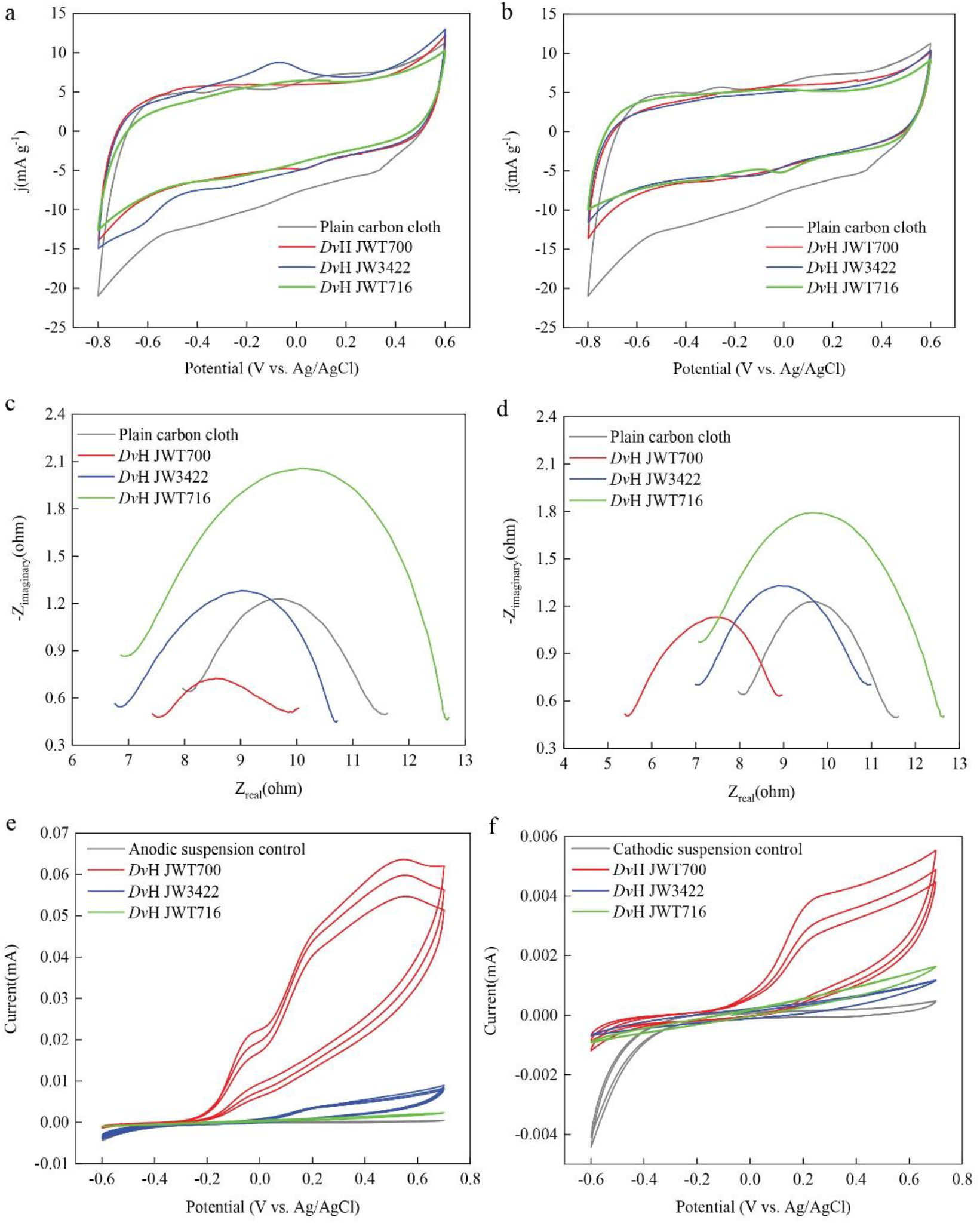
Electrochemical analysis of carbon cloths with different biofilms formed by various *Dv*H mutants. Cyclic Voltammograms for the anodes (a), cathodes (b), anodic media (e), and cathodic media (f) with different *Dv*H mutants. Nyquist impedance plots of the anodes (c) and cathodes (d) with different *Dv*H mutants’ biofilms.

The Nyquist plots analyzed by the classical equivalent electrical circuit are shown in Fig. 3 c and d. The diameter of semicircle represents the charge transfer resistance (R_ct_). The R_ct_ values of anodes with *Dv*H JWT700, *Dv*H JW3422, and *Dv*H JWT716 were found to be around 2.2, 3.7, and 5.7 Ω, respectively. This revealed that the biofilm formed by *Dv*H JW3422 was less conductive than the one formed by *Dv*H JWT700. Since anodes with DvH JWT700 biofilm had lower R_ct_ values than the plain carbon cloths (3.4 Ω), indicating that *Dv*H JWT700 formed the effective biofilm on the anode and promoted the electron transfer of electrodes. However, the cathodes with *Dv*H JWT700 and *Dv*H JW3422 biofilms had similar R_ct_ values with plain carbon cloths.

To investigate the components secreted by *Dv*H strains, further CV analysis was conducted for the anodic and cathodic suspension solutions. As a result, no redox peaks were found when a MOYLS4 medium without bacteria was used as the anolyte and catholyte (Fig. 3 e and f). Anodic suspensions with *Dv*H JWT700 and *Dv*H JW3422 in MOYLS4 had an oxidation peak in the forward scan of CVs at 0.45 V vs. Ag/AgCl. During the reverse scan, no reduction peak presents, which suggested oxidation activity because of the important contributions from *Dv*H JWT700 and *Dv*H JW3422. When the oxidation peaks of anodic suspension with *Dv*H JWT700 were much higher than that of *Dv*H JW3422, indicating that there were more oxidative components in the anodic suspension secreted by *Dv*H JWT700 than *Dv*H JW3422. Similarly, weaker peaks exhibited for a cathodic suspension of *Dv*H JW3422 and *Dv*H JWT716 compared to those of *Dv*H JWT700. It indicated that *Dv*H JWT700 secreted more oxidative components in the cathodic chamber. The weaker peaks redox by the cathodic suspension of *Dv*H JWT716 suggested that there might be redox components in the cathodic suspension that were secreted by the *Dv*H JWT716.

### 3.4 SEM and EDS analysis of different *Dv*H strains on electrode

Cells are curved rod-shaped (width of ∼0.5 μm and length of ∼3 μm) (Fig. 4). When growing on a mica sheet, *Dv*H primarily relies on its flagella to attach to the mica sheet surface (Fig. 4a and b). SEM images clearly revealed that biofilm formed by *Dv*H on the electrode surface, which has been retrieved from MFCs. It is worthy to note that *Dv*H biofilm formed on the surface of the anode contained a minimal amount of extracellular polymeric substance (EPS) materials that surrounded the cell (Fig. 4 c and d). In addition, only with filaments, *Dv*H also have pili with lengths of around 2 μm firmly attached to the cathode carbon cloth. The attachment was observed at the end of nano-filaments radiating out from a *Dv*H cell and sometimes between the attachment locations of adjacent filaments from *Dv*H cells. For comparison, *Dv*H JW3422 (non-pili forming one) was found containing EPS but without pili forming on the surface of the electrode (Fig. 4 e and f). Most *Dv*H JW3422 cells were attached to the surface of the cathode directly or through the EPS networks. As for the MFC cultured with *Dv*H JWT716, there were few cells attaching on the cathode (Fig. 4 g and h). EDS elemental spectra indicated enrichment of C, O and S in the deposits for *Dv*H JWT700 and *Dv*H JW3422 (Fig. S4).

**Figure 4.**
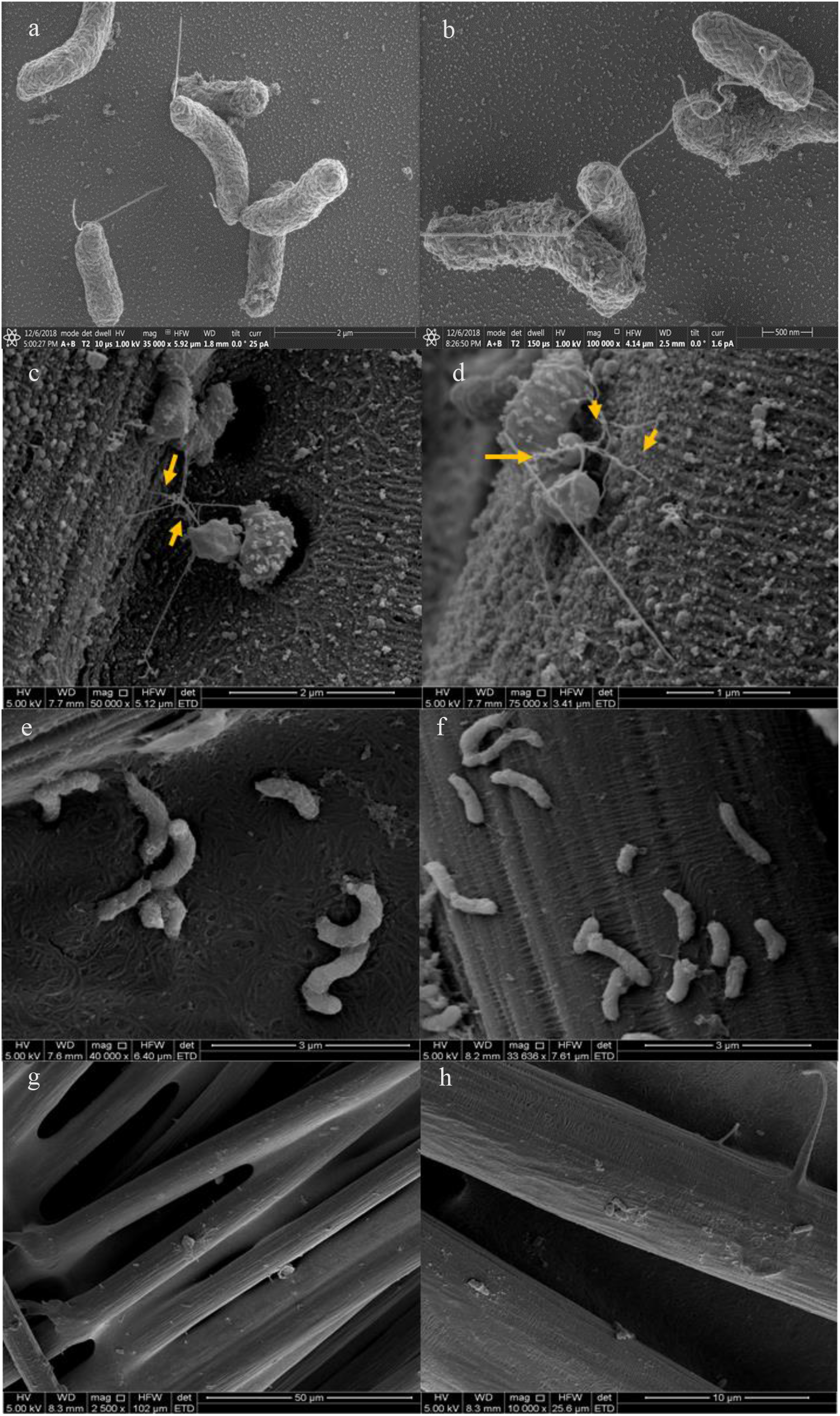
SEM micrographs of *Dv*H JWT700 (a∼d) growing on mica sheets, *Dv*H JWT700 (c∼d), *Dv*H JW3422 (e∼f) and *Dv*H JWT716 (g∼h) growing on carbon cloths. The yellow arrow represents the filaments formed by wild *Dv*H).

### 3.5 Electrically conductive pili of *Dv*H

The local structure of electrodes with *Dv*H biofilms and electronic properties were investigated through cAFM (Fig. S3). Topographic maps of the electrodes (carbon cloths) with biofilms were conducted by 2 µm × 2 µm to narrow down the location of microbe clusters and pili. The current was detected using a current-to-voltage preamplifier in the center topographic maps. Different voltages (i.e., 0∼2 V and 0∼4 V, respectively) were applied to plain carbon cloth and the carbon cloth with *Dv*H biofilm. The current signal of the carbon cloth with *Dv*H biofilm fluctuated greatly compared to the plain carbon cloth which had relatively flat signals (Fig. S3 b, d, f and h). Despite this, the conductivity of *Dv*H biofilm or pili cannot be determined since the carbon clothes are highly conductive.

### 3.6 Proteomics and metabolomics of electroactive respiration in *Dv*H

A total of 1149, 1400, 1208, and 1375 peptides were identified and found to be shared among different *Dv*H strains in four different electroactive respiration modes: on the anode (Fig. 5a), in the anodic suspension (Fig. 5b), on the cathode (Fig. 5c), and in the cathodic suspension (Fig. 5d), respectively. Upon comparing these peptides with the ones for biofilm and planktonic cultures of DvH JWT700 cultivated without electroactive respiration modes, it becomes evident that there is a higher percentage of unique peptides on the anode (comprising 16.4% of the total peptides found on the anode) and the cathode (accounting for 12.2% of the total peptides found on the cathode) when compared to their respective suspensions (constituting only 0.6% in the anodic suspension and 0.5% in the cathodic suspension). This suggests significant internal re-wiring in DvH to adapt to electroactive respiration compared to the dissimilatory sulfate reduction respiration of *Dv*H. Moreover, an increased presence of pili-related proteins (DVU2118, DVU1262, and DVU0451) and flagella-related proteins (DVU0513, DVU0311, DVU0514, and DVU0739) was noted for *Dv*H JWT700 and *Dv*H JW3422 on the anode. Similarly, a substantial presence of pili-(DVU2227, DVU1273, and DVUA0113) and flagella-(DVU0519, DVU0910, DVU3227, and DVU0045) related proteins was observed on the cathode for *Dv*H JWT700 and *Dv*H JW3422. These proteins including histidine kinase suggest an increased demand for surface attachment or motility. Furthermore, dominant unique peptides in the cathode for *Dv*H JWT700 and *Dv*H JW3422 include periplasmic [NiFe] hydrogenase (DVU2525) and [Fe] hydrogenase (DVU1771), underscoring their significance in the electroactive respiration process.

**Figure 5.**
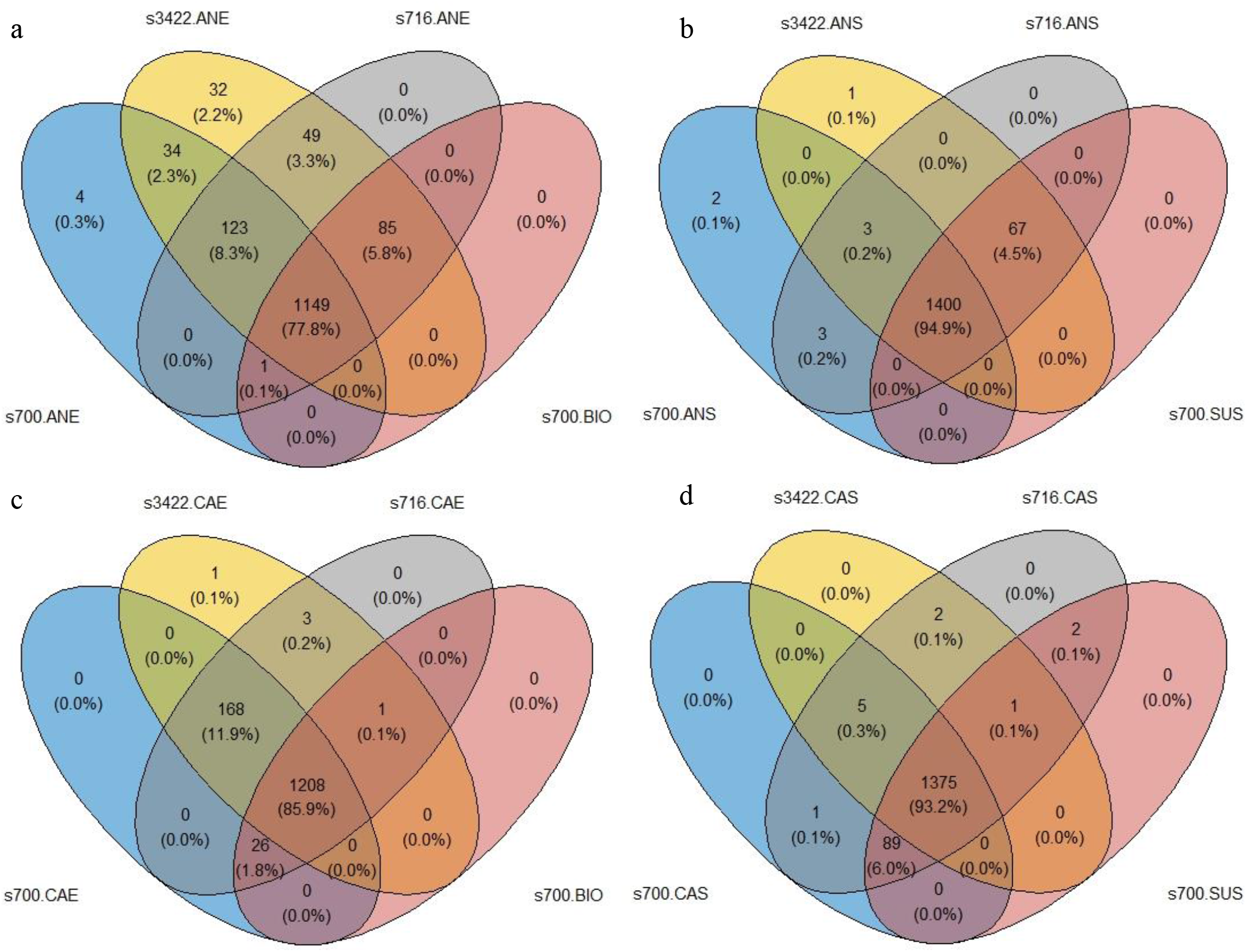
Venn diagram showing the number of unique peptides identified from *Dv*H JWT700 (s700; represented by the blue circle), *Dv*H JW3422 (s3422; represented by the yellow circle) and *Dv*H JWT716 (s716; represented by the grey circle) across various electroactive respiration modes. The labels ANE and ANS correspond to the anode and the suspension culture in the anodic chamber, respectively, while CAE and CAS represent the cathode and the suspension culture in the cathodic chamber, respectively. Additionally, s700.BIO and s700.SUS denote the biofilm and planktonic cultures of DvH JWT700 cultivated without electroactive repatriation modes.

For metabolites that were observed and validated in both positive and negative modes, we found riboflavin, flavin mononucleotide (FMN), and reduced flavin mononucleotide (FMNH_2_) appeared dysregulated between electroactive respiration modes (Table 5). By comparing it to the initial contraception riboflavin (5 ppm) in the media, we noticed that the riboflavin had been utilized with different levels. Riboflavin had been consumed in the anodic chamber culturing *Dv*H JW9003. FMN was observed in electroactive respiration modes of *Dv*H mutants. However, FMNH_2_ was produced by *Dv*H JWT700 and *Dv*H JW3422 under electroactive respiration, but not in the cathodic chamber culturing *Dv*H JWT716.

**Table 5.**
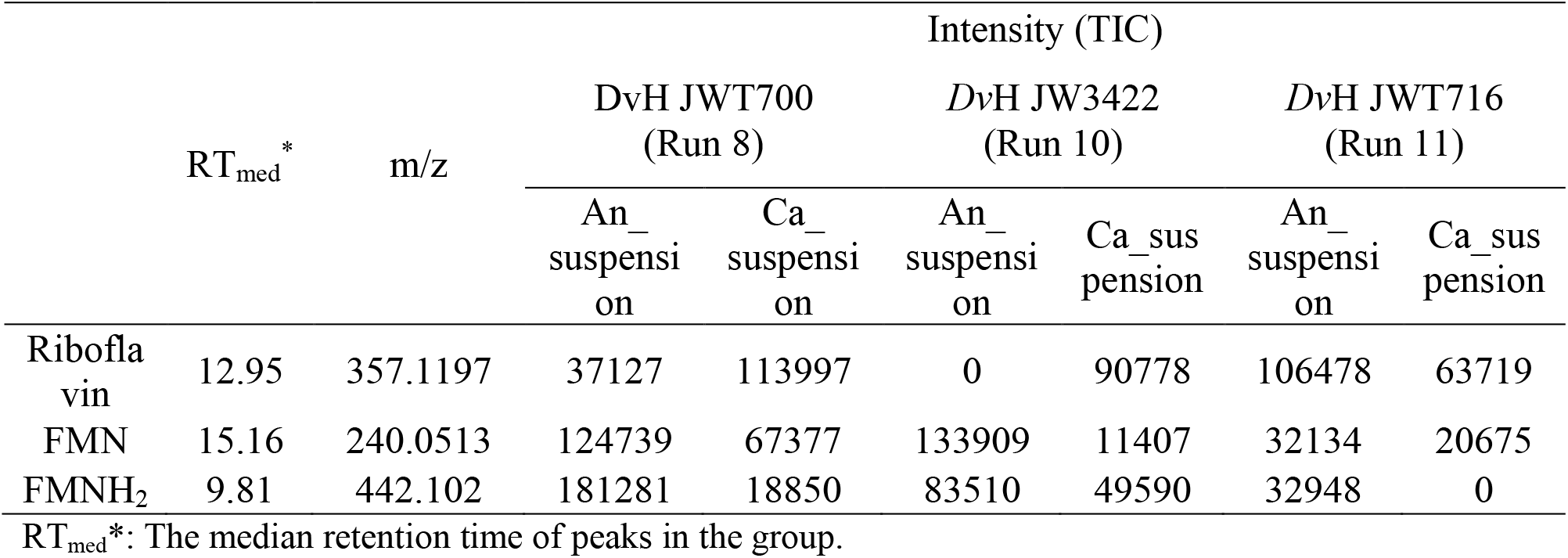
Flavin related metabolites of *Dv*H strains under different electroactive respiration modes.

|  | RT<sub>med</sub><sup>*</sup> | m/z | Intensity (TIC) |  |  |  |  |  |
| --- | --- | --- | --- | --- | --- | --- | --- | --- |
|  |  |  | DvH JWT700<br>(Run 8) |  | DvH JW3422<br>(Run 10) |  | DvH JWT716<br>(Run 11) |  |
|  |  |  | An_<br>suspensi<br>on | Ca_<br>suspensi<br>on | An_<br>suspensi<br>on | Ca_<br>suspensi<br>on | An_<br>suspensi<br>on | Ca_<br>suspensi<br>on |
| Riboflavin | 12.95 | 357.1197 | 37127 | 113997 | 0 | 90778 | 106478 | 63719 |
| FMN | 15.16 | 240.0513 | 124739 | 67377 | 133909 | 11407 | 32134 | 20675 |
| FMNH<sub>2</sub> | 9.81 | 442.102 | 181281 | 18850 | 83510 | 49590 | 32948 | 0 |

RT<sub>med</sub><sup>\*</sup>: The median retention time of peaks in the group.

## 4 Discussion

In this study, we observed that *Dv*H can harvest and send electrons to and from the electrode by varying the concentrations of the electron donor/electron acceptor ratio in the anodic chamber and cathodic chamber, respectively. By adopting different *Dv*H mutants (i.e., *Dv*H JW3422 and *Dv*H JWT716), we were able to determine the role of pili and the biofilm of *Dv*H. We also measured the metabolites under electroactive respiration for potential electron shuttles. The potential EET of *Dv*H has been proposed accordingly.

### 4.1 *Dv*H can send and receive electrons extracellularly from carbon cloth electrodes

With the supply of lactate in the anodic chamber, we applied the anode surface as the electron donor for the oxidizing reaction of lactate (runs 1-8). The cathode was employed as the electron donor for the reduction of sulfate. Fig. 2 showed that the variation of sulfate concentrations in the anodic chamber had more effects on the power generation than the variation of sulfate concentrations in the cathodic chamber. It has been reported that the COD/sulfate ratio has an influence on microbial activities in MFCs resulting in different potential outputs (48). Lee et al. found the MFC fed with the only sulfate yielded negligible electricity owing to lacking carbon source in a one-chamber, airbreathing cathode and continuous flow MFC at 22 °C (8). Our study has shown consistent results that when the sulfate concentration in the anodic chamber increased, electricity generation with the same condition in the cathodic chamber increased except for the ratio value equals to 2 (Fig. 2). By keeping the same condition of the anodic chamber, the increase of sulfate concentration in cathodic chamber resulted in an increase in electricity generation (Fig. 2b). One possible reason is that the high lactate/sulfate ratio approaching 2:1, which is the stoichiometric ratio of lactate and sulfate for *Dv*H (2C_3_H_5_O_3_^-^+ SO_4_^2−^+ 2H^+^ → 2C_2_H_3_O_2_^-^+H_2_S +2H_2_O + 2CO_2_), contributed to the optimal growth condition of *Dv*H (33) and led to more attachment of *Dv*H cells on the electrode. However, if the lactate/sulfate ratio equals two (Run 4), which led to the sulfate in the anodic chamber obtaining all electrons produced by lactate without transferring to the cathodic chamber. SBR that produce high coulombic efficiencies will lead to low biomass yields, as the electrons from the substrate are lost to produce current. In this study, most of the electrons from the lactate in the tests were employed to form biomass since our primary data showed that the increasing sulfate concentrations in the anodic chamber which may result in a higher biomass yield exhibited a higher coulombic efficiency. Additionally, the electricity generation from the MFC cultured *Dv*H indicated that *Dv*H cells may transfer electrons to the surface of the electrode and accept electrons extracellularly.

A higher power density was observed for *Dv*H in the MFC systems with a larger electrode size (Fig. 2c). Although *Dv*H JW3422 resulted in less power generation due to lack of nano-pili, they can still form the biofilm on the cathode surface (Figs. 2c and 3). It seems that the thin biofilm can also benefit the attachment of *Dv*H to the electrode surface and a nano-protein network structure was observed which may also help to transfer electrons from cells to cells and from cells to the electrode. It was previously demonstrated that extracellular polymeric substances include polysaccharides, proteins, glycoproteins, glycolipids, humic substances possess some semiconductive properties (49). According to the genome sequence information of *Dv*H, it produces vonWillebrand factor domain proteins, which are large, multimeric glycoproteins (50). These proteins could facilitate cell adhesion, pattern formation, and signal transduction, thus contributing to electron transfer. *Dv*H did not produce an extensive exopolysaccharide matrix and its biofilm formation dependent upon protein filaments (33). Different outer membrane *c*-type cytochromes such as MtrC, OmcA, OmcE, and OmcS offer various routes of electron transfer extracellularly for *S. oneidensis* and *G. sulfurreducens* (51, 52). Previous studies demonstrated that *Dv*H contains several membrane-bound redox complexes such as Qrc complex and Hmc complex that can accept electrons (53, 54). Qrc complex can accept electrons from the low-redox potential hemes of TpI*c*3 while Hmc transferred the electrons by transporting H^+^ from cytoplasmic lactate oxidation to the periplasmic cytochrome *c*_3_ network (55). However, none of these proteins were located at the outer membrane. Despite this, some uncharacterized proteins such as DVU1174, DVU0401, DVU1359, DVU0842 and DVU2997 on electrodes should be further investigated to determine their functions in EET of *Dv*H.

### 4.2 *Dv*H utilized filaments which facilitated electron transfer from cell-to-cell and to the electrode

CVs indicated the larger EDL capacitance of plain carbon cloths than the electrodes with *Dv*H biofilms (Fig. 3 a and b), which can be due to hydrogen sulfide produced by *Dv*H mutants poisoned the Pt wires on the surface of carbon cloths as shown the schematic in Figure 1 (56). Those poisoning largely reduced the pseudocapacitive contributions from Pt wires. A small redox peak in case of *Dv*H JW3422 revealed it might contribute to the oxidation reaction through unknown outer membrane *c*-type cytochromes. Furthermore, the lower R_ct_ of the anode with *Dv*H JWT700 demonstrated the formation of more effective biofilm on the anode that enhanced the electron transfer process (Fig. 3 d and c) as reported for *Klebsiella variicola* (57). However, non-pili forming *Dv*H JW3422 and non-biofilm forming *Dv*H JWT716 did not enhance the electron transfer process of anodes and non-biofilm forming *Dv*H JWT716 presented less conductive biofilm which may be due to the semiconductive EPS and the poisoning of Pt wires via hydrogen sulfide (56, 58). Irreversible oxidation processes appear in both the anodic and cathodic suspension of *Dv*H JWT700. In both chambers, the suspension of *Dv*H JWT700 had a larger oxidation current density than *Dv*H JW3422 and *Dv*H JWT716, suggesting *Dv*H JWT716 secreted more stable and oxidative compounds than the other two. These compounds might be quinone initiated or benzene derivatives chemicals or due to the similar CV curves presented in a previous study (59). In *Dv*H, flavin adenine dinucleotide (FAD) in quinone form which is the electron shuttle could accept two electrons and two protons to become hydroquinone form (FADH_2_) (60).

As discussed above, the pili seemed to play an important role in the electron transfer. We also noticed that nano-pili/filaments were effective for the attachment of cells of *Dv*H JWT700 on the surface electrode or functional as the networks between cells (Fig. 4). However, no such attachment using filaments was observed for *Dv*H JW3422. The biofilm formation of *Dv*H depends on protein filaments, which may mainly consist of pili and flagella. It was reported that the nano-pili produced by *Geobacter* genera were electrically conductive and they can facilitate electron transfers (52). Pili formed by *Dv*H was not fully investigated relating to the conductivity and structure. In this study, MFC cultivating non-pili forming *Dv*H JW3422 exhibited a lower power density (Fig. 2). Although the SEM-preparatory dehydration process may collapse or separate outer-membrane materials and break filaments, it was observed that *Dv*H exhibited little biofilm formation but produced numerous filaments or nano-pili on the surface of carbon cloths (Fig. 4). Additionally, pili-related proteins (DVU2118, DVU2227, DVU1262) were presented in the anodic chamber but were not found in *Dv*H under dissimilatory sulfate reduction respiration (Fig. 5 and Table S1). No definitive answer to the conductivities of *Dv*H’s pili can be obtained based on the current findings of cAFM.

Type IV pili can be categorized into two subclasses-type IVa pili and type IVb pili based on the sequence and length of the pilin subunit (61). *Geobacter sulfurreducens, Geobacter bremensis, Desulfuromonas thiophila*, and so on that were reported to have conductive pilin (e-pili) have type IVa structure (62). Although Holmes et al. (2016) reported pili in *Desulfovibrio vulgaris* were the long type IVa pilin, according to the previous finding of the sequencing alignment and 3D structure (Fig. S5), the *Dv*H pilin system is closer to the type IVb system. Until now, the pili structure of DVH is not well studied. Normally, the conductivity is depending on the composition of the amino acid chain of the major pili (63). A high density of aromatic amino acids and a lack of substantial aromatic-free gaps along the length of long pilins may be important to e-pilin (63). Two pilin proteins were found on the genome of *Dv*H: one is major pili, which belong to Flp family type IVb pili (DVU2116) (Fig. S6), and the other, prepilin-type N-terminal cleavage/methylation domain-containing protein, which may be the minor pili (putative PilE) of *Dv*H (Fig. S7). We further compared the major pilin and minor pili with the e-pili reported previously. No similar trend between sequences of the amino acid chain of major pili and e-pili was found. However, minor pili of *Dv*H had a lower E value and higher query cover percentage, indicating the minor pili shared a certain similarity with e-pili (Table S5). E-pili normally has phenylalanine (F), which aromatic amino acid, at the N terminus, and the majority have leader peptides with less than 12 amino acids (63). Instead of phenylalanine, the minor pili have Tyrosine (Y), which is also an aromatic amino acid. Thus, it is possible that the minor pili which are conductive may contribute to EET of *Dv*H. Being similar to *Dv*H, *D. desulfuricans* produces nanoscale filaments (32). These unidentified filaments were confirmed to be electrically conductive for extracellular electron transfer. *D. desulfuricans* also have both Flp family type IVb pilin and prepilin-type N-terminal cleavage/methylation domain-containing protein according to the identical protein group database in NCBI. Through the above analysis, pili of *Dv*H did not account for 100% of EET suggesting additional co-occurring direct or indirect mechanisms in this study.

Flagella of *Dv*H is composed of flagellin protein (i.e., DVU1441, DVU2444, and DVU2082) (64) and these proteins don’t have similarity with pilin of *Dv*H nor e-pilin. Flagellar and histidine kinase related proteins were dominant unique peptides in the anode and cathode comparing to dissimilatory sulfate reduction respiration (Fig. 5). It was reported that *Dv*H forms motility halos on solid media that are mediated by flagella-related mechanisms via the CheA3 histidine kinase (37). This indicated that *Dv*H had increased motility or surface attachment in the anodic chamber. In addition, previous studies suggested that biosynthesized FeS mediates the electron transport from *Dv*H to the electrode surface (31, 65). For instance, Deng et al. (2020) found *Dv*H biosynthesized FeS nanoparticles on the cell membrane in the presence of sulfate and iron as an electron conduit enabling *Dv*H to utilize solid-state electron donors via direct electron uptake (65). However, no biosynthesized FeS was observed on the surface of the cathode based on the results of EDS elements analysis, but they were observed as precipitates at the bottom of MFCs. The FeS nanocrystallites may be washed off during the fixation preparation process of SEM.

### 4.3 *Dv*H uses electron shuttles for indirect electron transfer

*Shewanella* species was found to secreted flavins (i.e., FMN and riboflavin) as electron shuttles to bind to outer membrane cytochromes mediating EET (66). Although *Geobacter* species have abundant *c*-type cytochromes and are thought to transfer electrons by direct contact, flavin synthesis and excretion genes are widely distributed in *Geobacter* species (67). Studies indicated that *Geobacter sulfurreducens* can uptake self-secreted riboflavin as bound cofactors for EET (68). Since we have provided riboflavin in the media, it is not surprising that we detected the riboflavin through the metabolite analysis (Table 5). However, under electroactive respiration, *Dv*H strains have different levels of utilization on riboflavin. Riboflavin is considered the precursor of flavin nucleotides (i.e., FMN and FAD). During the catalytic cycle, FMN cycles between FMN to reduced flavin mononucleotide (FMNH_2_) enable the electron transfer via redox reactions. *S. oneidensis* MR-1 can use the interaction of flavin/outer membrane *c*-type cytochrome complexes to regulate the extracellular electron transport (69). Compared to *Dv*H under dissimilatory sulfate reduction respiration, more riboflavin synthase (e.g., DVU1199, DVU1200 and DVU1201) have been detected in anodic suspension under electroactive respiration. It was previously found both riboflavin and FAD accelerated pitting corrosion and weight loss on the stainless steel caused by *Desulfovibrio vulgaris* biofilm (67). Thus, FMN and FMNH_2_ may act as electron shuttles of *Dv*H. However, the cell-surface redox-active proteins need to be investigated to determine the free-flavin-mediated electron-shuttling mechanism of *Dv*H.

### 4.4 DvH can employ a combination of electron transfer mechanisms to use solid surfaces as both an electron acceptor and a donor

The reported coulombic efficiency in the current study is much lower (Tables 2 and 3) than the ones reported in other relevant studies, which varied from 6.7% to 98.9% when different feed compositions and SRB were applied (8, 70, 71). The highest current densities come from mixed cultures that are usually dominated by the genus *Geobacter* (72). The differences in coulombic efficiency were caused by different MFC configurations and SRB species, which have different electron transfer mechanisms or use different electron donors (17). To date, no comprehensive information on the EET mechanism of *Dv*H from cells to electrode was exhibited. DET based on FeS clusters, conductive pili/filaments, unknown outer membrane *c*-type cytochromes, and EPS, and IET based on electron shuttles were potential EET for *Dv*H but not completely understood (Fig. 6). Besides the DET conducted by FeS nanoparticles (73), our results indicated that direct contact through pili/filaments may be one of the major routes for DET of *Dv*H. Pili/ filaments also contribute to the attachment of *Dv*H to the surface of electrodes. In a previous study, extracellular enzymes, such as hydrogenase and formate dehydrogenase could mediate a direct electron uptake for *Methnococcus maripaludis* (74). However, *Desulfovibrio* species mainly have cytoplasm-located and periplasmic hydrogenases which contribute to the intracellular EET and hydrogen formation (75, 76). More outer membrane redox complexes are needed to be investigated to reveal the possibility of DET dominated by these proteins. Our results showed that IET mechanisms via electron shuttle small molecules which are flavin-related ones are also happening in *Dv*H. This indicates that *Dv*H can use a combination of electron transfer mechanisms to survive in less than ideal conditions with solid surfaces as both an electron acceptor and a donor. A previous study reported that the presence of nanowires of *Candidatus Desulfofervidus* HotSeep-1 cell depended on substrates-hydrogen, which contribute to the interspecies electron transfer (77). However, whether *Dv*H can secrete riboflavin, FMN and FMNH_2,_ the relevant transporters through the membrane and the contributions in extracellular electron transfer need to be further determined. It has been reported that *Deltaproteobacteria* have unique and unknown metabolisms and *Dv*H belongs to this group, indicating *Dv*H might also have some metabolic abilities involving dissimilatory sulfate reduction respiration under various conditions. The alignments also showed that bacteria in *Deltaproteobacteria* may have similar functions or ecological roles without having the most similar genomes. This might indicate *Dv*H may have similar electron transfer mechanisms with other bacteria in *Deltaproteobacteria*, such as *Geobacter* sp.

**Figure 6.**
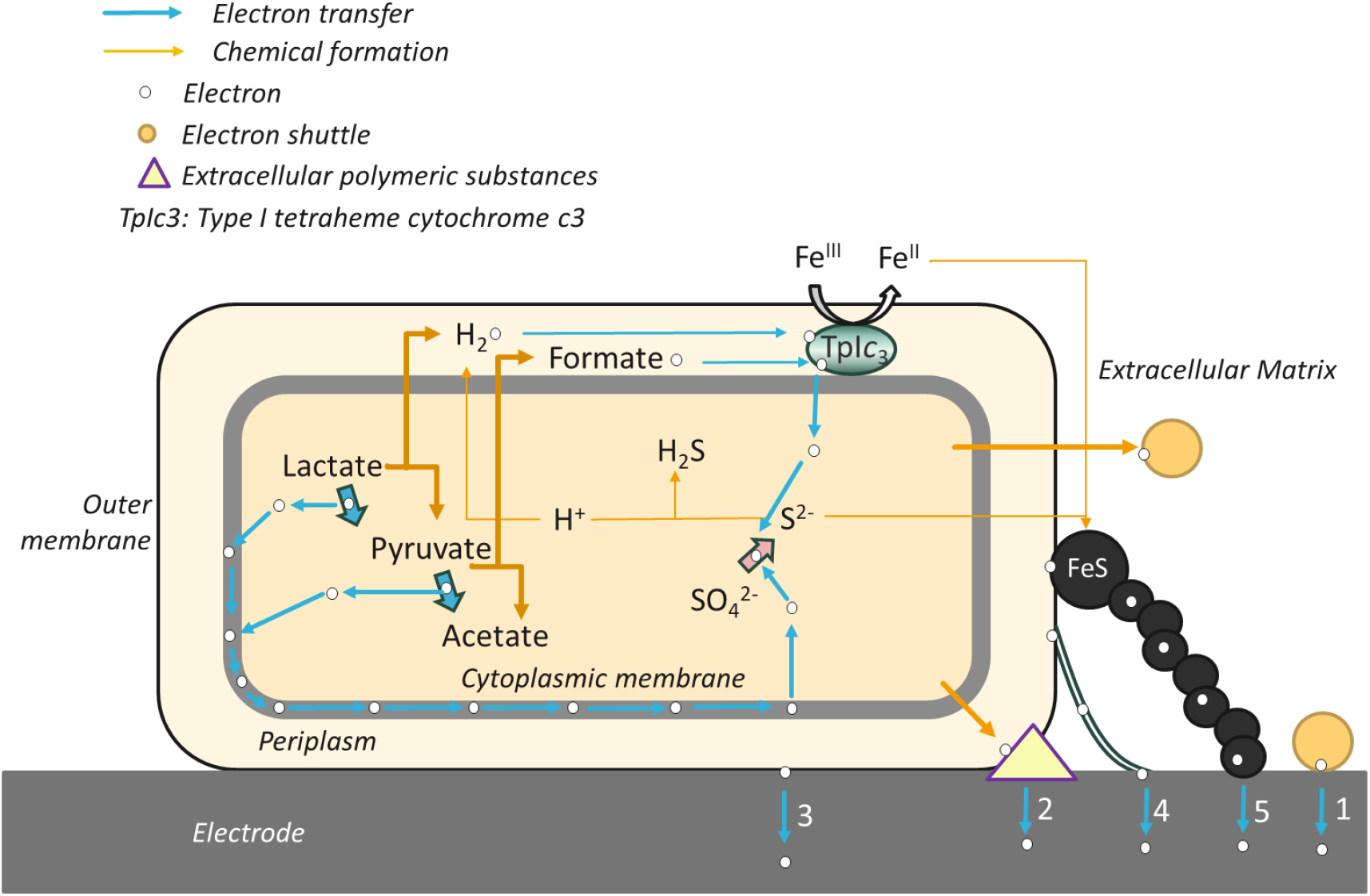
All potential EET Mechanisms of *Dv*H. IET based on electron shuttles (1) and DET based on EPS (2), unknown outer membrane *c*-type cytochromes (3), conductive pili/filaments (4), and FeS clusters (5) are presented.

Previous studies focused mainly on the biocorrosive capacity of *Dv*H (35, 78) and overlooked the electricity production capacity of *Dv*H through MFC. *Dv*H employs a combination of mechanisms, including both direct and indirect contributions, to achieve electroactive respiration. A comprehensive understanding of *Dv*H’s electron transfer mechanisms can expand its application beyond MFCs to address sulfate-containing water and wastewater in various contexts and applications.

## 5 Conclusions

In this study, we found that *Dv*H was able to attach onto the electrodes of MFC, resulting in the formation of nano-filaments on the electrode surface and electricity production with a maximum power density of ∼0.074 W/m^2^. Varying the sulfate concentration in the anodic chamber appeared to have more effects on electricity generation than that in the cathodic chamber with *Dv*H enriched in both chambers. *Dv*H was found to utilize nano-filaments/pili to attach the cells on the electrode surface and facilitate electron transfer from cell to cell and to the electrode. In contrast, an MFC inoculated with the *Dv*H mutant with a deletion of the major pilus-producing gene was found to generate less voltage and exhibit reduced attachment to the electrode surface forming fewer biofilms. Untargeted metabolomics profiling showed flavin-based metabolites, potential electron shuttles. This indicated that *Dv*H has possibly multiple direct electron transfer pathways to grow on solid surfaces as the electron acceptor and donor. Future work is needed to confirm the composition of pili and apply the enhancement of pili in situ to overcome MFC inefficiencies and generate more current.

## Acknowledgements

The authors acknowledge funding from the University of Missouri Discovery Challenge Award and the Excellence in Electron Microscopy Award, SUNY Research Foundation, and Wisconsin Alumni Research Foundation. We would especially like to thank Dr. Judy D. Wall for the support, strains and lab space during the initial work on this project. Thanks are given to: Grant Zane and Dr. Kara B De Leon for providing *Dv*H mutants; Deanna Grant for SEM sample preparation; Dr. Ebbing de Jong at SUNY Upstate Medical Center for initial global proteomics analysis and Grzegorz ‘Greg’ Sabat from the Mass Spectrometry Core Facility in the Biotechnology Center at the University of Wisconsin-Madison for performing proteomics; Dr. Qiang Gao for assisting with CV analysis; Dr. Andrew Hryckowian for lending anaerobic chambers and Dr. Alexander B. Artyukhin for LC-MS method set-up.

